# Natural selection promotes the evolution of recombination 2: during the *process* of natural selection*

**DOI:** 10.1101/2021.06.07.447324

**Authors:** Philip J Gerrish, Fernando Cordero, Benjamin Galeota-Sprung, Alexandre Colato, Varun Vejalla, Nick Hengartner, Paul Sniegowski

**Affiliations:** Department of Biology, University of New Mexico, Albuquerque, New Mexico, USA; Department of Biology, University of Michigan, East Lansing, Michigan, USA; Theoretical Biology & Biophysics, Los Alamos National Lab, Los Alamos, New Mexico, USA; Department of Mathematics, Universidad Autónoma del Estado de Hidalgo, México; Biomathematics and Theoretical Bioinformatics, Technische Fakultät, Universität Bielefeld, Germany; Department of Biology, University of Pennsylvania, Philadelphia, Pennsylvania, USA; Departamento de Ciências da Natureza, Matemática e Educação, Univ Fed de São Carlos, Araras SP, Brazil; Thomas Jefferson High School for Science and Technology, Alexandria, Virginia, USA

## Abstract

The ubiquity of sex and recombination in nature has eluded unified explanation since the time of Darwin. Conditions that promote the evolution of recombination, broadly defined as any form of genetic mixing, are fairly well understood: it is favored when genomes tend to contain more selectively mismatched combinations of alleles than can be explained by chance alone. Yet, while a variety of theoretical approaches have been put forth to explain why such conditions would prevail in natural populations, each has turned out to be of limited scope and applicability. Here, we show, simply and surprisingly, that natural selection acting on standing heritable variation always creates conditions favoring the evolution of recombination, in expectation. Specifically, we find that, in expectation: 1) the mean selective advantage of recombinants is non-negative, 2) the mean selective advantage of a recombination-competent modifier is non-negative, and 3) the asymptotic frequency of a recombination-competent modifier is close to one and is independent of the strength of selection. Remarkably, these findings are independent of the distribution of genic fitnesses in the standing heritable variation upon which natural selection acts, implying that the source of this variation is immaterial. Taken together, our findings indicate that: 1) the evolution of recombination should be promoted in expectation wherever natural selection is operating, and 2) sex and recombination may have evolved more as a byproduct than as a catalyst of natural selection.

## I. INTRODUCTION

The oldest ideas about the evolutionary role of recombination are from Weismann (1889), who argued that sex provides increased variation for natural selection to act upon. Since then, the amount of work that has addressed the evolution of sex and recombination is spectacular. To preface our developments therefore, we cover some essential background and make reference to some reviews [3–9] that give a much more complete overview of the remarkable wealth of previous and current work in this area. Also, we refer to our companion publications ev0 [1] and ev1 [2] for additional introductory material.

Fisher [10] and Muller [11] first provided concrete mechanisms for an advantage to recombination. Muller surmised that in order for separately arising beneficial mutations to fix in the same genotype, in an asexual population they must arise in the same lineage sequentially, while in a recombining population, they may arise contemporaneously and be subsequently reshuffled into the same background. Fisher argued that a single beneficial mutation, because it arises in a single individual, has a significant probability of arising on a non-optimal genetic background. In an asexual population, the beneficial mutation is stuck with this non-optimal background, while in a recombining population, the background can be swapped out for a fitter one. If the beneficial mutation is successful, despite arising on a non-optimal background, a second beneficial mutation may eventually arise on the background of the first as it progresses toward fixation. If this happens in an asexual population, Hill and Robertson [12, 13] found that the probability of success of the second beneficial mutation will be depressed as a consequence of arising in the growing lineage founded by the first beneficial mutation. Generally speaking, genetic linkage (the absence of recombination) introduces selective interference [5] that decreases the efficiency of natural selection.

Recombination can ameliorate all of these linkage-induced hindrances to natural selection [14–17], and recombining populations should therefore adapt faster [18]. However, the magnitude of this benefit depends very much upon parameter choices [19]. More fundamentally, this process provides only a group-level benefit for sex, and group-level explanations, besides being characteristically viewed with suspicion in evolutionary biology, are unsatisfactory in that they cannot explain the origin and fixation of sexual reproduction within a single population, nor explain its maintenance by evolution [20]. Therefore it is necessary to study the evolution of recombination within a single population.

To do so requires consideration of an additional “modifier” gene (or *locus*), variants of which (or *alleles*) determine recombination rate. An allele at this locus conferring increased recombination rate is introduced into a population at low frequency. The questions of interest are: 1) what is the selective value of this allele? And 2) what is the fate of this allele? A variety of theoretical studies have modeled the evolution of such recombination modifiers. These studies have investigated mechanisms including fluctuating selection [14, 15, 21]; negative epistasis [14, 15, 22, 23]; assortative mating [24]; and finite population effects, i.e. drift [12, 17, 25–27]. Of these, the drift-based explanations have come into favor in recent years as the more promising in explaining the ubiquity of recombination [28], but the general consensus is that some fundamental piece of the puzzle is still missing [5].

A modifier can itself be subject to the very recombination it modulates and can thus have limited-term linkage to the fitness loci whose recombination rate it modifies. Whether the selective value of recombination is determined by short-term or long-term effects depends on how long a modifier will typically remain linked to the fitness loci whose recombination rate it modifies; a loosely-linked modifier will be affected by short-term effects whereas a tightly-linked modifier will be affected by both short- and long-term effects. We derive the selective value and dynamics of a recombination-competent (*rec*^+^) modifier under loose and tight modifier linkage.

To address the evolution of sex and recombination, we have taken a reductionist approach. Our aim is restricted to studying the effects of one very key process, namely natural selection, in isolation (no mutation, no drift, etc), and we distill this problem to what we believe is its most essential form: we ask, how does the action of natural selection, by itself, affect the selective value and fate of recombinants and recombination? In choosing this approach, we sought analytic tractability and transparency that might lead to robust new insights into the evolution of sex and recombination.

In companion papers ev0 [1] and ev1 [2], we show that natural selection acting on standing variation has an encompassing tendency to amplify selectively mismatched combinations of alleles, thereby promoting the evolution of recombination across the *products* of selection, defined as genotypes that have become locally prevalent in their respective populations, subpopulations, demes, or clones through the local action of natural selection. In the present study, we assess how the selective value of recombinants and recombination are affected during the *process* of natural selection within an unstructured population. In these combined studies, we find that recombinants are favored and recombination promoted, in expectation, as an inherent consequence of the dynamics and statistical properties of selective sorting.

## II. FITNESS EVOLUTION

As stated above, the goal of the present study is to focus exclusively on natural selection and ask how natural selection, by itself, affects the selective value of recombinants and recombination. This goal requires a reductionist approach in which natural selection is studied in isolation. Consequently, our evolutionary models here retain only the natural selection terms; other, more complete models that incorporate selection, mutation, drift, and recombination, may be found in the SM; these are presented there for completeness and to lay the groundwork for subsequent studies.

### One locus

This model is a continuous-time formulation of evolution by natural selection; the model and its analyses are not new and have close parallels in [29–34]. We let *u*_*t*_(*x*) denote probability density in fitness *x* at time *t* (i.e., ∫_*x*_ *u*_*t*_(*x*) = 1 for all *t*) for an evolving population. Dropping the subscript *t*, we have that, under selection and mutation, *u* evolves as:

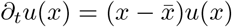

where 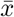 is mean fitness 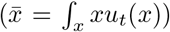. Let *M* (*φ*) denote the moment-generating function (*mgf*) for *u*(*x*), i.e., 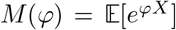, where random variable *X* has density *u*(*x*). The transformed equation is:

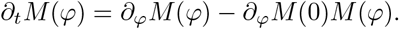

We define cumulant-generating function (*cgf*) *𝒞*(*φ*) = ln *M* (*φ*); noting that *∂*_*φ*_ *𝒞* (*φ*) = (*∂*_*φ*_*M* (*φ*))*/M* (*φ*), and *∂*_*t*_𝒞(*φ*) = (*∂*_*t*_*M* (*φ*))*/M* (*φ*) we find that the *cgf* evolves as:

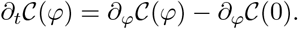

This equation is a variant of the transport equation; it is immediately apparent that the solution will be of the form *𝒞*_*t*_(*φ*) = *F* (*φ* + *t*), where *F* is an arbitrary function. When boundary condition *𝒞*_*t*_(0) = 0 ∀ *t* is applied, it has solution:

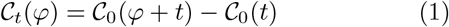

where the subscripts are now necessary again: *𝒞*_*t*_(*φ*) is the *cgf* of the fitness distribution *u*_*t*_(*x*) at time *t*. We note that the fitness evolution of a population can thus be projected into the future based only on the present fitness distribution (i.e., at *t* = 0).

### Two loci

While many parts of our analyses are true for general fitness functions, where total fitness is some arbitrary function *z* = *ϕ*(*x, y*), we here and in other parts of our analyses restrict ourselves to additive fitness *z* = *x* + *y*. Remarkably, however, this restriction to additive fitness will cease to be necessary in some expressions and key findings downstream from here. Also, generalizations of these evolution equations to non-additive functions are found in the SM.

We now suppose that there are two “genes” that determine fitness. Letting fitness contributions of the two genes be denoted by *x* and *y*, respectively, the total fitness is *z* = *x* + *y*. The extension of the previous one-dimensional *pde* is immediate:

Let *u*_*t*_ (*x, y*) denote probability density in fitness contributions *x* and *y* at time *t* for an evolving population. Dropping the subscripts again, under selection and mutation, *u* evolves as:

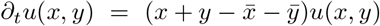

The *cgf* is now two-dimensional: 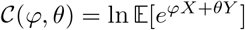, where random variables *X* and *Y* have density *u*(*x, y*). The *cgf* evolves as:

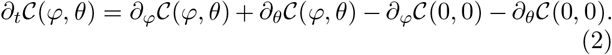

This equation is a two-dimensional variant of the transport equation and has more possible solution forms than the one-dimensional case, namely, solutions can be of the form: *F* (*t* + *φ, θ − φ*), *F* (*t* + *θ, φ − θ*), or *F* (*t* + *φ, t* + *θ*). The consistent solution form is the last of these. When boundary condition *𝒞*_*t*_(0, 0) = 0 ∀ *t* is applied, it has solution:

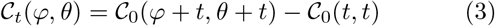

where the subscripts have again become necessary. We again note that the evolution of a population can thus be projected into the future based only on the present fitness distribution (i.e., at *t* = 0).

### Finitely-many genotypes

The foregoing developments are expressed in terms of analytical *cgf* ‘s, which implicitly assumes that a population is of infinite size and consists of an infinite number of genotypes. Real populations contain a finite, possibly even small, number of genotypes. The foregoing developments are equally applicable when number of genotypes is finite, in which case the analytical *cgf* ‘s are replaced with empirical *cgf* ‘s defined as follows:

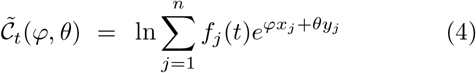

where *x*_*j*_ and *y*_*j*_ are genic fitnesses of genotype *j, n* is number of genotypes, and *f*_*j*_(*t*) is the frequency of genotype *j* at time *t*. In what follows, *f*_*j*_ written as such with no argument denotes initial frequency, i.e., *f*_*j*_ := *f*_*j*_(0). In many of the developments that follow, we will assume that genotypes are initially found at equal frequency so that *f*_*j*_ = 1*/n* for *j* = 1, 2, …, *n*.

### Evolution with recombination

Let *u*(*x*, ·) denote the marginal density of individuals carrying genic fitness *x* at the first locus, and let *u*(·, *y*) denote the marginal density of individuals carrying genic fitness *y* at the second locus. Independent evolution at the two loci means that *u*(*x, y*) = *u*(*x*, ·)*u*(·, *y*). Let 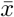 denote the mean of 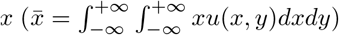. Under recombination, we have the two loci evolving independently as follows:

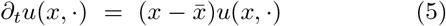

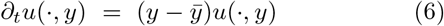

In the previous section, we have shown that transformed versions of the above evolution equations have the closed-form analytical solutions:

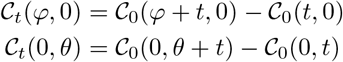

under free recombination. Mean fitness of the *rec*^+^ sub-population at time *t* is therefore:

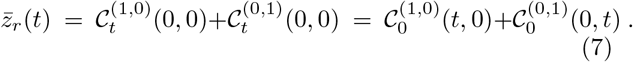

### Evolution without recombination

For the non-recombining wildtype subpopulation, the evolution equation has already been presented and analyzed above. The transformed equation has solution:

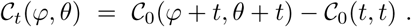

Mean fitness at time *t* is now:

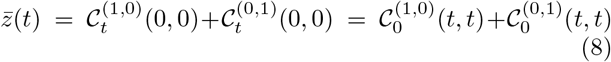

### Evolution of fitness differential between *rec*^+^ modifier and wildtype

The fitness differential between the *rec*^+^ modifier and wildtype at time *t* is:

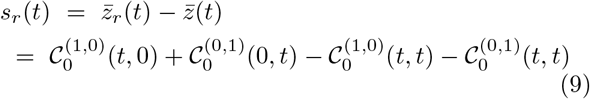

where 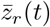 and 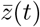 are given by Eqs (7) and (8). This may be equivalently written as:

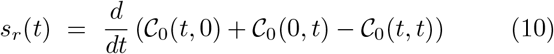

or as

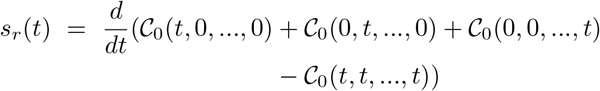

for more than two loci.

We note that future dynamics of the fitness differential is predicted based simply on the distribution of fitnesses in the initial variation upon which natural selection acts, i.e., *s*_*r*_(*t*) depends only on *𝒞*_0_(*φ, θ*), the *cgf* of the initial fitness distribution.

### Asymptotic fitness differential between *rec*^+^ modifier and wildtype

#### Proposition 1.

*A population initially consists of n distinct genotypes at equal frequency (this assumption is relaxed later) characterized by their genic fitness vector* (*x*_*i*1_, *x*_*i*2_, …, *x*_*im*_), *i* = 1, 2, …, *n. These values may be drawn from any multivariate distribution, continuous or not. The action of natural selection by itself will cause the fitness of a* rec^+^ *modifier to increase relative to its non-recombining counterpart by an amount given by:*

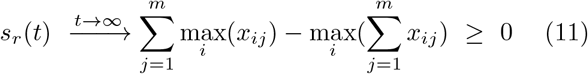

*Proof*: We employ the fitness differential as defined by Eq (9) and insert empirical *cgf* ‘s as defined by Eq (4), giving:

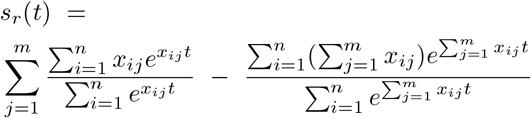

which gives the result by inspection. Inspection is facilitated by examining the case of two loci whose genic fitnesses we deonte *x*_*i*_ and *y*_*i*_:

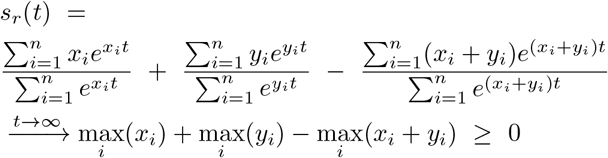

□

#### Corollary 1.

*For the case of two genotypes and two loci (n* = 2 *and m* = 2*) the asymptotic fitness differential given by Proposition 1 can be rewritten as:*

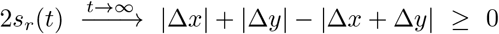

where Δ*x* = *x*_2_ − *x*_1_ and Δ*y* = *y*_2_ − *y*_1_.

Remark: The couples (*x*_1_, *y*_1_) and (*x*_2_, *y*_2_) are two independent draws from some unspecified bivariate distribution. This fact guarantees that Δ*x* and Δ*y* are symmetric, from which it is apparent that the asymptotic fitness differential will be zero half of the time.

#### Corollary 2.

*We now define random variable S*_*r*_ *to denote the asymptotic fitness differential as defined above. Here we generalize Proposition 1, representing fitness at each of the m loci by random variables X*_*j*_, *j* = 1, 2, …, *m. Expected asymptotic fitness differential is:*

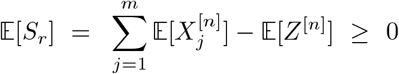

*where* 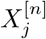 *denotes the n*^*th*^ *order statistic (i*.*e*., *maxima) of X*_*j*_, *j denotes the j*^*th*^ *locus, and Z*^[*n*]^ *denotes the n*^*th*^ *order statistic (i*.*e*., *maxima) of* 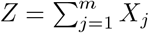.

The foregoing expressions very accurately predict the fitness differential after one bout of selection in simulations (Fig 1), and are surprisingly robust to recombination rate and initial *rec*^+^ frequency.

**FIG. 1.**
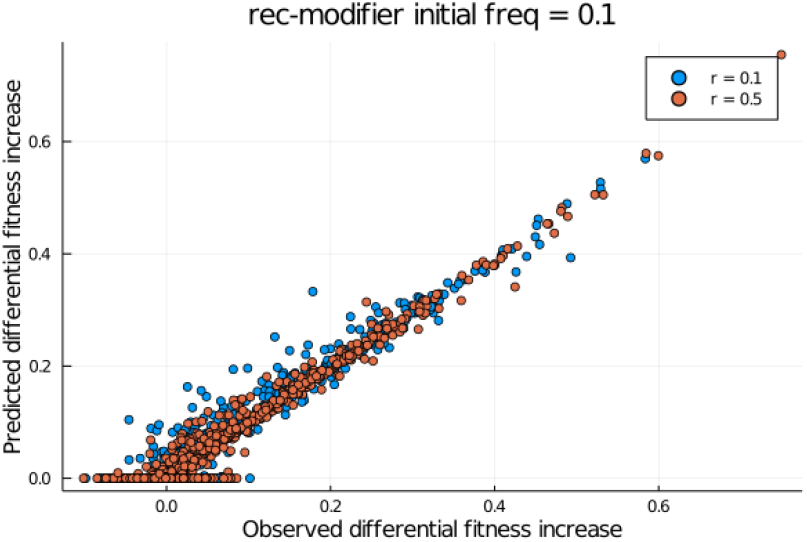
Fitness differential between *rec*^+^ modifier and wild-type after one bout of selection: observed (horizontal axis) and predicted (vertical axis). Initial modifier frequency was 0.1, and plotted are values for recombination rates of *r* = 0.1 (blue) and *r* = 0.5 (red). Observed values come from fully-stochastic simulations. Predicted values are computed using Eq (11)

### Dynamics of a *rec*^+^ modifier

To understand how natural selection affects the *evolution* of recombination, the more directly relevant question to ask is how natural selection affects the *frequency* of a *rec*^+^ modifier.

If a lineage has time-dependent selective advantage *s*(*t*), its frequency *ρ*(*t*) evolves as the solution to the logistic equation:

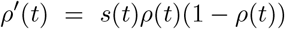

which, with initial condition *ρ*(0) = *ρ*_0_, has solution:

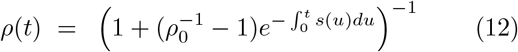

At this point, our choice of *s*(*t*) will reflect the nature of the *rec*^+^ modifier in question and, in particular, whether the modifier is loosely or tightly linked to the fitness loci whose recombination rate it modifies. As we will show, both cases derive from Eq (10): for the loose-linkage case, *s*(*t*) = *ŝ*_*r*_(*t*) as defined below; for the tight-linkage case, *s*(*t*) = *s*_*r*_(*t*).

### *Loose linkage modeled as* rec^+^ *dominance*

In our simulations, loose linkage is modeled by considering the *rec*^+^ allele at the modifier locus to be dominant, following Felsenstein [35]. A dominant *rec*^+^ modifier, when paired with *rec*^−^ wildtype, will result in a phenotypically *rec*^+^ pair of coupled genomes. While technically speaking, diploidy is required for dominance, our use of haploid dominance is substantively indistinguishable from diploid dominance for our purposes. The phenotypic *rec*^+^ expression of *rec*^+^/*rec*^−^ pairs means that the modifier can be dissociated from the fitness loci whose recombination rate is modified. In this case, what determines the dynamics and fate of the *rec*^+^ modifier is the short-term gain conferred by the modified recombination rate, i.e., the short-term selective value of recombinants. A recessive *rec*^+^ modifier, on the other hand, models tight (or complete) linkage between the modifier and fitness loci, because only *rec*^+^/*rec*^+^ pairs will have the *rec*^+^ phenotype. Exchange at the modifier locus in this case is irrelevant because a *rec*^+^ allele can only be exchanged for another *rec*^+^ allele.

## III. LOOSELY-LINKED *REC* ^+^ MODIFIER

### Loose linkage gives most conservative case

When a modifier is loosely linked to fitness loci, the fate of the modifier is only weakly determined by the selective value of the recombinants it forms. Loose linkage thus embodies the most conservative case in terms of the magnitude of natural selection’s effect on recombination. Tight linkage, while increasing the magnitude of the effect of natural selection, may nevertheless go in the opposite direction of that determined for the loose-linkage case. This possibility of opposite effects is implied in Barton’s identification of a distinction between short- and long-term effects [14]. In the present manuscript, we investigate both of these cases and find that, when natural selection acts in isolation, the effects under loose and tight linkage are in the same direction, namely favoring recombination.

### Negative covariance: the measure of *rec*^+^ advantage under loose linkage

Incomplete linkage is formulated by letting 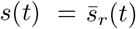, defined as follows. We let *τ* denote the characteristic duration of linkage between modifier and fitness loci. Tighter linkage will result in larger *τ*. Under incomplete linkage, the mean selective advantage of a modifier at time *t* derives directly from Eq (10) and is given by:

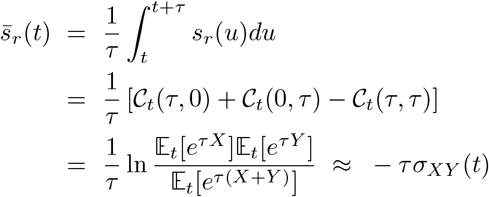

The approximation made by the last step is very accurate for small *τ* (see SM) because the two-dimensional Jensen gaps for numerator and denominator essentially cancel each other out. In other words, under incomplete linkage, the selective advantage of a *rec*^+^ modifier is −*τσ*_*XY*_ (*t*). As *τ* grows, we find numerically that these developments provide a conservative bound: 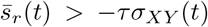 (see SM).

Because we are interested primarily in the sign of the selective value of recombination, we can let *τ* = 1 without loss of generality. To make our language precise, we define “loose linkage” to mean *τ* = 1, and we let *ŝ*_*r*_ denote the selective value of recombination under loose linkage; we note that the selective value of recombination is equal to the selective value of recombinants in this case. The term “incomplete linkage” will refer to *τ >* 1, and we let 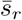 denote the selective value of recombination under incomplete linkage (as employed above).

### Covariance dynamics

Covariance dynamics can be forecast given only the distribution of genic fitnesses at time zero; this derives immediately from Eq (3):

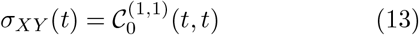

where 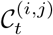(*φ, θ*) denotes the (*i, j*)th derivative of the *cgf* for genic fitnesses at time *t*.

#### Infinitely-many genotypes

We note that if the initial fitness distribution is normal, covariance remains constant over time: *σ*_*XY*_ (*t*) = *σ*_*XY*_ (0) ∀ *t*.

In real populations, the number of genotypes is finite. Given the marked qualitative difference between the finite- and infinite-genotypes cases, we will focus primarily on the finite-genotypes case.

#### Finitely-many genotypes

Replacing *x*_*j*_ and *y*_*j*_ in Eq (4) with random variables *X*_*j*_ and *Y*_*j*_, the future dynamics of expected covariance may be computed as:

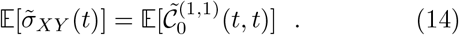

This covariance-forecasting equation is shown to be very accurate in Fig 2.

**FIG. 2.**
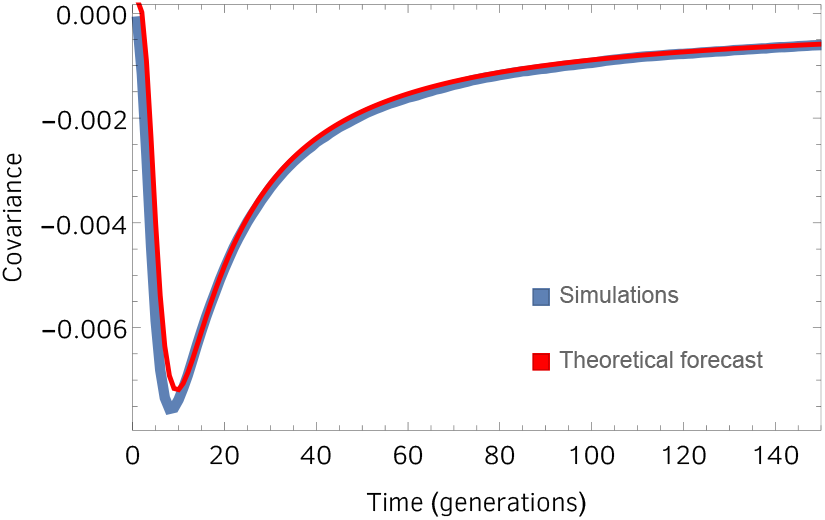
Covariance forecast. Blue curve is the average of 1000 stochastic, individual-based simulations. Red curve is the theoretical prediction given by 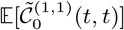 (Eq (14)). We call it a forecast because it is based solely on the distribution of fitnesses at *t* = 0. Fitnesses were drawn from a bivariate normal distribuiton with means (0, 0) and standard deviations (0.2, 0.2) and zero correlation. The initial population consisted of ten distinct genotypes in a population of size *N* = 2000.

### Without epistasis

We recall that recombinant advantage is −*σ*_*XY*_. Here, we examine the simplest scenario of two loci and two genotypes. We study how the selection-driven changes in the frequencies of types (*x*_1_, *y*_1_) and (*x*_2_, *y*_2_) *within a single unstructured population* change covariance *σ*_*XY*_ = *σ*_*XY*_ (*t*) over time. We are interested in the net effect of these changes, given by 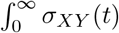; in particular, we are interested in knowing whether this quantity is positive (net recombinant disadvantage) or negative (net recombinant advantage) in expectation.

#### Proposition 2.

*Within-population covariance integrated over time is:*

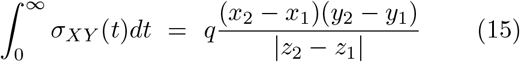

*where q is the initial frequency of the inferior genotype. And z*_*i*_ = *x*_*i*_ + *y*_*i*_.

*Proof*: We employ Eq (14) to give us covariance dynamics as a function of the initial two genotypes. We let *p* denote initial frequency of the superior of the two genotypes, and we let *q* = 1 − *p* denote initial frequency of the inferior genotype. Time-integrated covariance is:

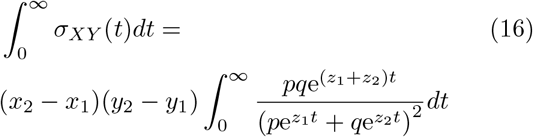

We note the convenient fact that the integral depends only on the *z*_*i*_, allowing us to relax the assumption of additive fitness and revert to the general definition *z*_*i*_ = *ϕ*(*x*_*i*_, *y*_*i*_) – a fact we will leverage below. Integration by parts yields:

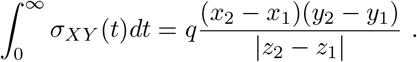

□

Now to find the expectation of time-integrated covariance, we must replace *x*_*i*_ and *y*_*i*_ with random variables *X*_*i*_ and *Y*_*i*_.

#### Proposition 3.

*We define random variables* Δ*X* = *X*_2_ − *X*_1_, Δ*Y* = *Y*_2_ − *Y*_1_, *and* Δ*Z* = *Z*_2_ − *Z*_1_ = Δ*X* +Δ*Y*. *The* (*X*_*i*_, *Y*_*i*_) *are independently drawn from any distribution (with any correlation), making* Δ*X and* Δ*Y centered and symmetric. Time-integrated covariance is unconditionally non-positive in expectation:*

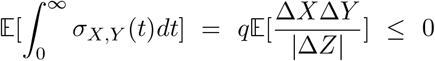

*This is equivalent to saying that time-averaged selective advantage of recombinants is unconditionally non-negative*.

*Proof*: The bivariate distribution governing *X* and *Y* is irrelevant to this proof: no assumption about this distribution is required. There is also no need to assume that (Δ*X*, Δ*Y*) has a density. Δ*X* and Δ*Y* are two real-valued random variables such that: (− Δ*X*, Δ*Y*) has the same distribution as (Δ*X*, Δ*Y*). This is guaranteed by the fact that Δ*X* and Δ*Y* are spacings. We have:

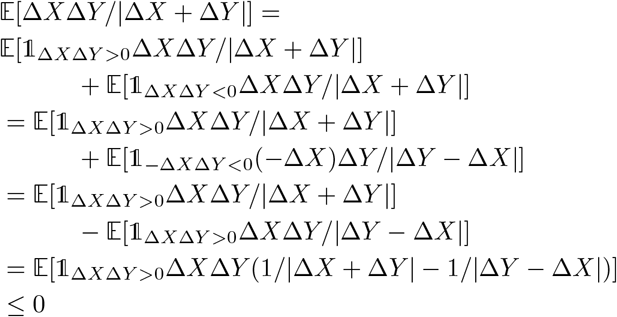

When Δ*X* and Δ*Y* have the same sign as imposed by the indicator function in the last expectation, we have |Δ*X* + Δ*Y* |*>* |Δ*Y* − Δ*X* |, from which the inequality derives. □

#### Corollary 3.

*Proposition 3 holds for divergent expectations*.

*Proof*: Set *U* = |Δ*X*| and *V* = |Δ*Y*| ; Λ = Max(*U, V*), *λ* = Min(*U, V*). Then you can rewrite the expectation as:

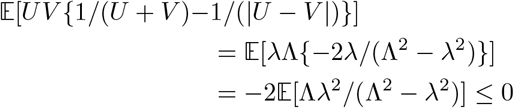

Indeed, if the expectation is divergent, then it is always −∞. This approach removes the need to make the argument that *U* + *V >* |*U* − *V*| and avoids the need to take a difference of expectations. An alternative approach is given in an expanded statement and proof of Proposition 3 in the SM. □

### With epistasis

We now leverage the aforementioned fact that fitness additivity is not required here and we employ the general definition of *z*, namely *z* = *ϕ*(*x, y*) which we insert into Eq (16), leading to the following corollary to Proposition 3:

#### Corollary 4.

*For any real number ξ, let us consider a fitness function of the form ϕ*_*ξ*_(*x, y*) = *aX* + *bY* + *ξg*(*X, Y*), *where a, b >* 0 *and g is a function independent of ξ. Let* Δ*Z*(*ξ*) = *ϕ*_*ξ*_(*X*_2_, *Y*_2_) − *ϕ*_*ξ*_(*X*_1_, *Y*_1_). *Assume that for some ε >* 0,

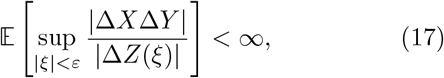

*and that* ℙ(Δ*X*Δ*Y* = 0) *<* 1. *Then, there is ε*_0_ ∈ (0, *ε*), *such that for all ξ* ∈ (−*ε*_0_, *ε*_0_), *we have*

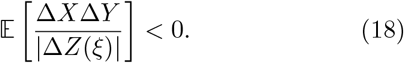

*Proof*. Condition (17) implies that the function *h* : (−*ε, ε*) → ℝ defined via

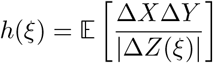

is continuous. Moreover, since ℙ(Δ*X*Δ*Y* = 0) *<* 1, proceeding as in the proof of Proposition 3, we obtain that *h*(0) *<* 0. Hence, by continuity of *h*, we infer that there is *ε*_0_ ∈ (0, *ε*) such that *h* is negative in (− *ε*_0_, *ε*_0_), which concludes the proof.

Figure 3 plots the left-hand side of the inequality in Eq (18) with generalized fitness function:

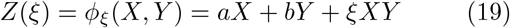

with *a, b >* 0 and epistasis parameter *ξ ∈* ℝ. This figure reveals where the interval (− *ε*_0_, *ε*_0_) lies for different correlation coefficients. The predicted symmetry of this interval about zero is corroborated with both Montecarlo expectations of the left-hand side of Eq (18) as well as fully-stochastic evolutionary simulations.

**FIG. 3.**
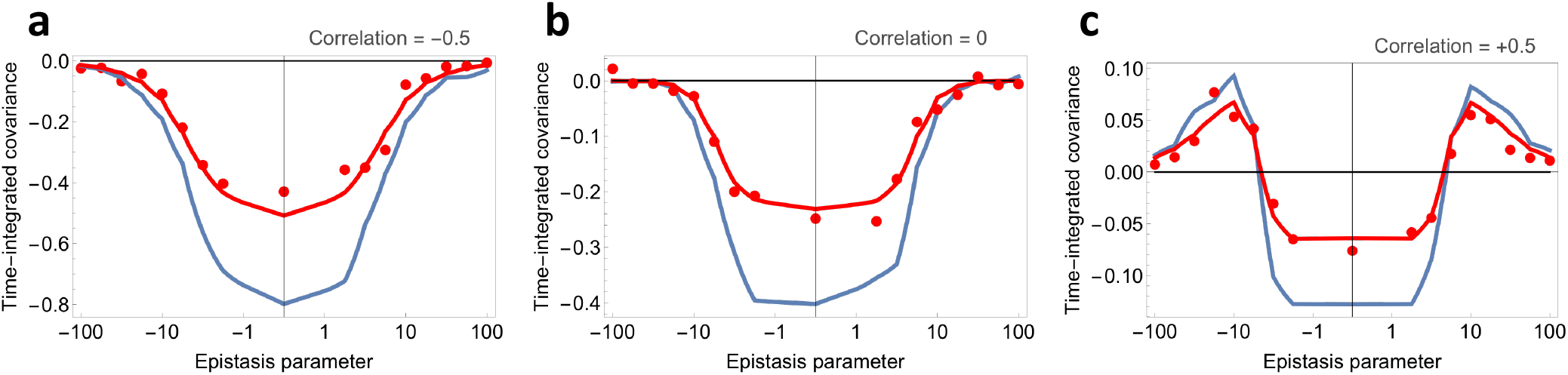
Effect of epistasis *ξ* (horizontal axes) on time-integrated covariance (vertical axes) when correlation between genic fitnesses in the initial population is: **a)** − 0.5, **b)** 0, and **c)** +0.5. Theoretical predictions (solid curves) plot Montecarlo expectations 𝔼 [Δ*X*Δ*Y/* Δ*Z*(*ξ*) |], the left-hand side of Eq (18), where *N* = 2000 (red) and *N* = 10^5^ (blue). Red points plot the means of 500 stochastic simulations with *N* = 2000. For these plots, we employ the general fitness function given by Eq (19). These plots support the validity of our conjecture that *Z* in Eq (16) can be generalized by allowing *Z* = *ϕ*(*X, Y*). They also corroborate our analyses showing that time-integrated covariance is unconditionally negative in the absence of epistasis. They further corroborate our finding that time-integrated covariance is negative in an epistasis interval that is symmetric about zero. Finally, they indicate that epistasis must be fairly strong (either positive or negative) to make time-integrated covariance non-negative.

We now turn our attention to the analysis of time-integrated covariance with epistasis for the special case where total fitness *Z* is given by Eq (19). As before, we let Δ*Z*(*ξ*) = *ϕ*_*ξ*_(*X*_2_, *Y*_2_) − *ϕ*_*ξ*_(*X*_1_, *Y*_1_) = (*a* + *ξY*_1_)Δ*X* + (*b* + *ξX*_2_)Δ*Y*. The case where the random variables (|Δ*X*Δ*Y*| */* |Δ*Z*(*ξ*) |)_*ξ*∈(−*ε,ε*)_ are uniformly integrable (i.e. condition (17) is satisfied) is covered already by Corollary 4. If it is not uniformly integrable, we have the following:

#### Corollary 5.

*Assume that the distribution of* (*X*_*i*_, *Y*_*i*_) *has finite support, i*.*e. there is K >* 0 *such that* ℙ(*X*_*i*_ ∈ [ − *K, K*], *Y*_*i*_ ∈ [ − *K, K*]) = 1 *and that* |*ξ*| *<* (*a* ∧ *b*)*/K, where a* ∧ *b denotes the minimum between a and b. The we have:*

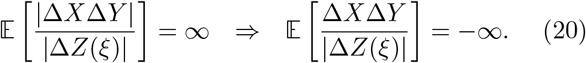

*The proof for this corollary is in the SM*.

### Modifier dynamics under incomplete linkage

In foregoing developments, we have shown that the selective advantage of a *rec*^+^ modifier under incomplete linkage is:

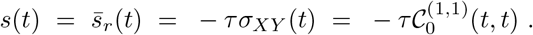

Furthermore, we have shown that:

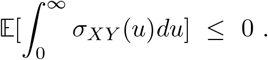

Inserting the above expression for *s*(*t*) into Eq (12), and employing Jensen’s inequality, we have:

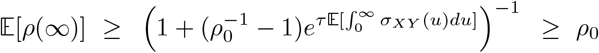

as long as 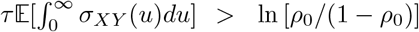, which defines the region for which *ρ*^′′^(*t*) *>* 0; this condition will hold quite generally for any biologically reasonable properties of the *X* and *Y*. Figure 4 plots asymptotic modifier frequency, *ρ*(∞), for the case of *n* = 10 distinct genotypes. This figure shows that, under incomplete linkage, while the average increase in modifier frequency is not large, it is never negative.

**FIG. 4.**
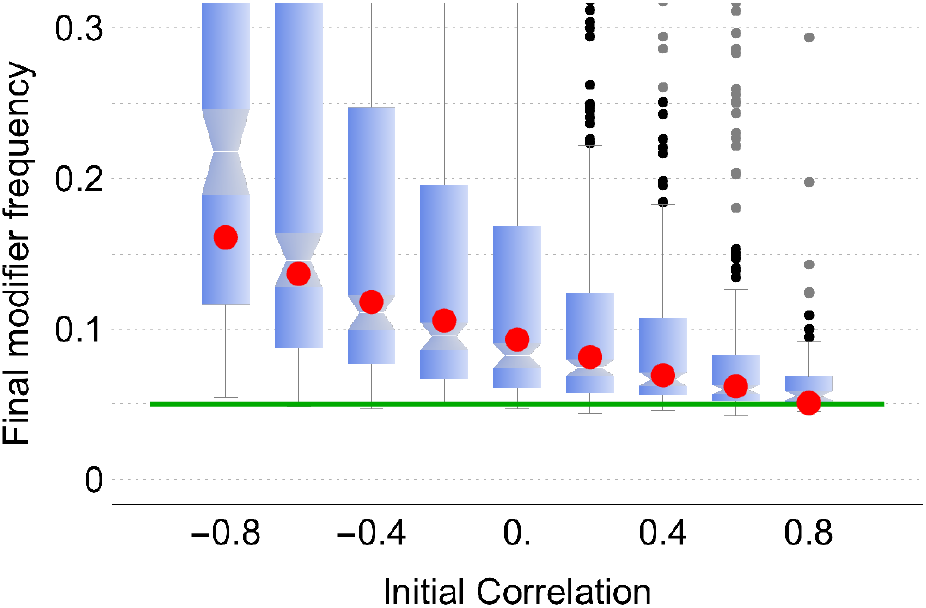
Asymptotic modifier frequency under incomplete linkage. Initial modifier frequency was 0.05, indicted by the green line. The box-whisker chart plots Montecarlo realizations of the analytical expression for *ρ*(*∞*). Red dots plot the means of 5000 simulations of natural selection acting in populations of size 2000. Initially, populations consisted of 10 distinct genotypes at equal frequency. Each genotype was constructed by drawing genic fitness values *X* and *Y* from a bivariate normal distribution with zero means, variances equal to 0.04 and correlation specified by the horizontal axis. Each simulation was run until fixation occurred. Analytical asymptotic modifier frequency is given by the expression 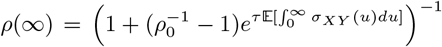. Parameter *τ* is the characteristic duration of linkage between modifier and fitness loci and was determined to be *τ ≈* 2.1. Black and gray dots plot outliers. Significantly, there are no outliers below the initial modifier frequency of 0.05.

In sum, a dominant *rec*^+^ modifier will not decrease in frequency (on average) as a consequence of natural selection, despite being only loosely linked to fitness loci. This effect increases (*rec*^+^ modifier grows to higher frequencies) when more genotypes and/or loci are considered.

## IV. TIGHTLY-LINKED *REC* ^+^ MODIFIER

Conveniently, we immediately have the anti-derivative of *s*_*r*_(*t*) from Eq (10) from which we have the definite integral:

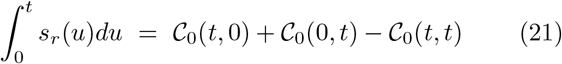

We replace 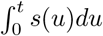 in Eq (12) with 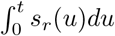 from Eq (21), giving the expression for modifier dynamics:

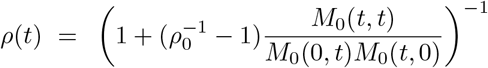

This expression gives the generalized case for two loci and holds for any number *n* of genotypes. The further generalization to *m* loci and *n* genotypes is immediate:

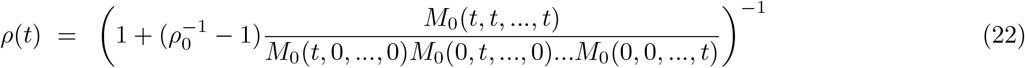

### Infinitely-many alleles

Assuming there are infinitely-many alleles, and supposing that the distribution of *X* and *Y* is bivariate normal, we have:

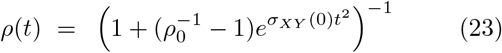

so modifier dynamics depend critically on the sign of the covariance of the initial fitnesses: when *σ*_*XY*_ (0) *<* 0, then 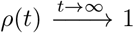; when *σ*_XY_ (0) *>* 0, then 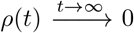; and when *σ*_*XY*_ (0) = 0, then *ρ*(*t*) = *f* (0) ∀ *t*. This finding stands in stark contrast to the case of finitely-many alleles in which, as we shall see, modifier frequency for all practical purposes always ends up at higher frequency than where it started, quite independently of the bivariate distribution governing *X* and *Y* (even when the initial population has strongly positive correlation between *X* and *Y*).

Remark: In general, Eq (23) may be written for any distribution:

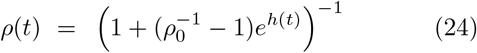

where:

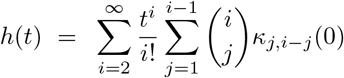

and *κ*_*i,j*_(0) is the (*i, j*)^*th*^ cumulant of the initial bivariate distribution of genic fitnesses *X* and *Y*. This expression is obtained by Taylor expansion of Eq (21). The normal case can be gleaned from this general expression by recalling that for the normal distribution, *κ*_*i,j*_(0) = 0 when *i* + *j >* 2, so that *h*(*t*) = *κ*_1,1_(0)*t*^2^ = *σ*_*XY*_ (0)*t*^2^.

### Finitely-many alleles

If the number of alleles is finite, we employ the empirical *cgf*, 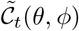, as defined by Eq (4), and the empirical *mgf*, 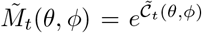. Assuming *n* genotypes are present in the population in question, and replacing *𝒞* with 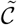 in Eq (21), we have:

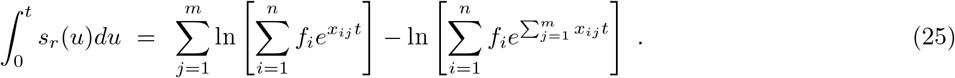

Assuming for now that *f*_*i*_ = 1*/n*, the expected frequency of the modifier is:

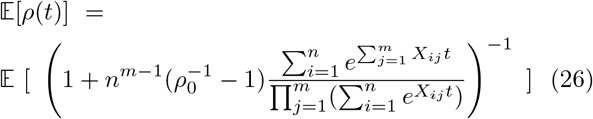

From here on, we will restrict ourselves to the case of *m* = 2 loci for the sake of presentation. The general *m*-locus case is a rather trivial (albeit messy) extension of these developments. For the two-locus case, Eq (26) becomes:

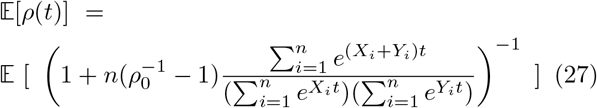

which can be rewritten as:

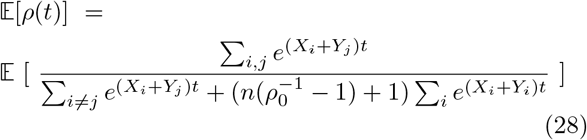

Equation (27) accurately predicts *rec*^+^ modifier dynamics as show in Fig 5.

**FIG. 5.**
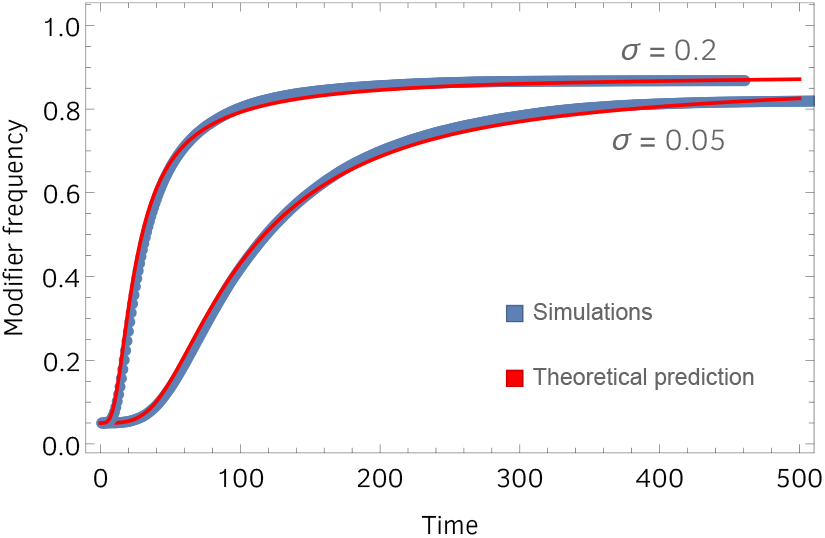
Recombination modifier dynamics over the course of one bout of selection. Red curves plot theoretical predictions given by Eq (27). Blue curves plot mean trajectory observed in 500 replicate simulations. Simulations were fully stochastic, individual-based, with a population size of *N* = 20, 000. The recombination-competent modifier conferred a recombination rate of *r* = 0.2. The distribution of genic fitnesses *X* and *Y* in the initial population had a bivariate normal distribution with zero means, standard deviations *σ*_*X*_ = *σ*_*Y*_ = *σ* = 0.2 for the upper curves and *σ*_*X*_ = *σ*_*Y*_ = *σ* = 0.05 for the lower curves, and zero correlation. The initial population consisted of *n* = 10 distinct genotypes. We note that upper and lower curves eventually converge to the same asymptotic frequency, as our theory predicts, despite very different strengths of selection (very different *σ*’s).

We note that, for the case *n* = 2, Eq (27) has the curious alternative form:

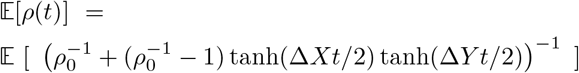

where Δ*X* = *X*_2_ − *X*_1_ and Δ*Y* = *Y*_2_ − *Y*_1_.

### Asymptotic modifier frequency

In foregoing developments, we have shown that, after one “bout” of selection has completed, the fitness advantage of a tightly-linked *rec*^+^ modifier is non-negative. This is indeed suggestive of natural selection’s favorable effect on such a modifier, but it only gives information about the modifier’s fitness advantage at the *end* of the bout of selection. It does not guarantee, for example, that the modifier’s fitness advantage did not become negative over the course of the bout of selection and consequently suppress the modifier’s frequency in the process. This concern is especially relevant in light of our observation in the remark following Corollary 1 that, for the case of two loci and two alleles, the modifier’s fitness advantage after the bout of selection has completed is zero half of the time.

#### Proposition 4.

*A non-recombining population initially consists of n distinct genotypes. The i*^*th*^ *genotype is characterized by the random vector of genic fitnesses* (*X*_*i*1_, *X*_*i*2_, …, *X*_*im*_), *where m is the number of loci under selection. This random vector may have any multivariate distribution, continuous or not. A* rec^+^ *modifier is introduced into the population at frequency ρ*_0_. *The action of natural selection by itself will cause the frequency of the modifier to converge in expectation to:*

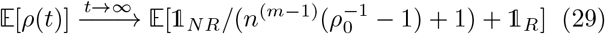

*where conditions NR and R are met when the maximum-fitness genotype is a non-recombinant and recombinant, respectively. Specifically, NR is met when the maximum-fitness genotype has the following property: for any pair of genic fitnesses* (*X*_*ij*_, *X*_*kl*_), *it is the case that i* = *k. Condition R is met when NR is not true*.

*Proof*:

The proof is by inspection of Eq (28) and its full *m*-locus extrapolation. (Inspection is facilitated by first considering the case *m* = 2.) □

From here, it is easy to see that *in theory* the modifier can decrease in frequency to below its initial frequency. This happens under the worst-case scenario for the modifier, which is when the correlation coefficient becomes extremely (unrealistically) close to +1. When the correlation is exactly equal to +1, we have:

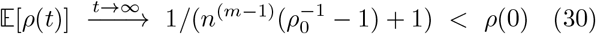

where *n* ≥ 2 is the number of genotypes in the initial population and *m* the number of loci constituting a genotype. Numerical solution of Eq (29), however, reveals that the correlation coefficient has to be unrealistically close to one for the modifier to decrease in frequency (Fig 6).

**FIG. 6.**
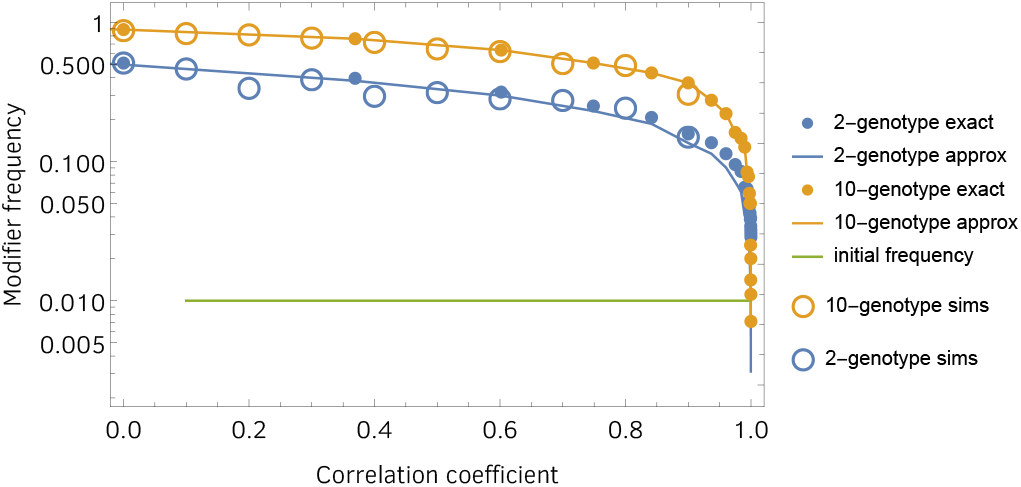
Asymptotic modifier frequency as a function of positive correlation between genic fitnesses *X* and *Y* among the genotypes initially present in the population, as computed by Eq (29). Only when the correlation gets unrealistically close to one does the asymptotic modifier frequency dive to values that can, in theory, dip below the initial frequency. Asymptotic modifier frequency for negative correlations is not plotted; it becomes increasingly close to one as correlation becomes increasingly negative. Open circles plot final modifier frequency in simulated populations of size 2000 with recombination rate of the modifier of *r* = 0.1.

#### Corollary 6.

*We generalize Proposition 4 by allowing each genotype i to have its own starting frequency f*_*i*_ 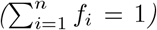. *Expected asymptotic modifier frequency in this generalized case is:*

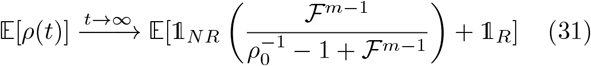

*where conditions NR and R are as defined above, and random variable ℱ is starting frequency (the f*_*i*_ *are instances of ℱ)*.

If *ℱ* is distributed as one dimension of an *n*-dimensional flat Dirichlet distribution (ensuring 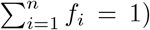, and *m* = 2, Eqs (29) and (31) are, for all practical purposes, equivalent. For *m >* 2, we have found numerically that:

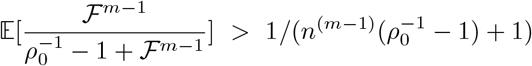

but the left-hand side is still very small, validating the following Corollary for equal or random starting frequencies.

#### Corollary 7.

*If a* rec^+^ *modifier is initially at low frequency in a population, it’s final (asymptotic) frequency is well approximated by the probability that the maximum-fitness genotype is a virtual recombinant*.

*More specifically, given that a population initially consists of n genotypes carrying random vectors of genic fitnesses* (*X*_*i*1_, *X*_*i*2_, …, *X*_*im*_), *i* = 1, 2, …, *n, the final expected modifier frequency is effectively equal to the probability of condition R defined in Proposition 4 above*.

*Proof*:

This corollary comes about by noting that the first term on the right-hand side of Eq (29) is typically much smaller than the second term:

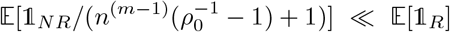

for the case of equal starting frequencies, or:

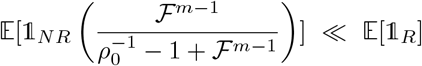

for the case of random starting frequencies. This fact is corroborated by Montecarlo expectations plotted in Fig 6 where this approximation appears indistinguishable from the exact solution. □

Corollary 7 allows us to say some things about non-extreme cases as well. When the initial covariance is zero, Corollary 7 together with simple combinatorics tell us the asymptotic frequency is:

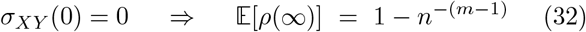

If the heritable variation upon which natural selection acts is itself a product of previous selection, our companion papers [1, 2] show that the initial covariance will be negative. For this case, asymptotic modifier frequency is even higher:

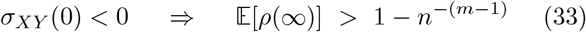

from which it is apparent that asymptotic modifier frequency quickly gets very close to one as number of genotypes and number of loci increase. For example, two genotypes and ten loci will have an asymptotic modifier frequency greater than 0.998 when the correlation is negative. Significantly, Eqs (32) and (33) appear to be fairly robust to epistasis (SM).

A surprising feature of the asymptotic modifier frequency is the absence of any requirement for information about the magnitude of selective differences among alleles at the different loci. This independence of strength of selection is illustrated in Fig. 5, where it is corroborated with simulations. An implication of this finding is that modifier frequency will converge to the same value even under weak selection. This fact may speak to concerns, expressed in previous work [7, 36], about the strength of selection required for recombination (and sex) to evolve.

And perhaps even more surprising is the fact, shown in Fig 6, that increase in modifier frequency is substantial even when the correlation between *X* and *Y* is strongly positive in the initial population. It is only when this correlation gets unrealistically close to +1 that increase in modifier frequency is substantially reduced. This is surprising because a strongly positive correlation between genic fitness is precisely the condition that one would expect to suppress, not favor, recombination (discussed in [1, 2]). The reason that modifier frequency increases despite positive correlation has to do with the dynamics of selective sorting, and the fact that these dynamics cause recombinants to be favored, on average (covariance to be negative on average), independently of the correlation between genic fitnesses *X* and *Y* in the initial population or any other feature of those fitnesses for that matter, as proved in Proposition 3 above.

## V. EVOLUTIONARY DYNAMICS WITH RECOMBINATION

Let *u*_*t*_(*x, y*) denote probability density in fitness contributions *x* and *y* at time *t* for an evolving population. Dropping the subscripts, under selection and recombination, *u* evolves as:

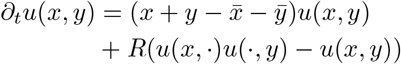

where *R* is recombination rate. The transformed equation is:

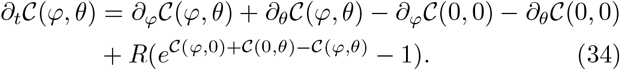

whose solution is given by the *𝒞*_*t*_(*φ, θ*) which satisfies:

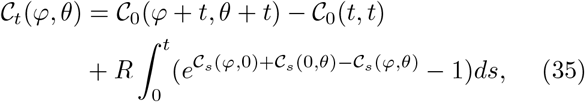

and boundary condition, *𝒞*_*t*_(0, 0) = 0 ∀ *t*. This equation can be solved iteratively for *𝒞*_*t*_(*φ, θ*).

These developments lead to a variant of Proposition 2:

### Proposition 5.

*A first iteration of* *Eq* (35) *yields a modification of the time-integrated within-population covariance given in Proposition 2:*

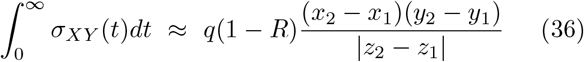

*where q is the initial frequency of the inferior genotype, and continuous parameter R* ∈ [0, 1] *is recombination rate*.

*Proof*: For the first iteration, we simply replace the *𝒞*_*s*_(*φ, θ*) in the exponent by *𝒞*_0_(*φ* + *s, θ* + *s*) −*𝒞*_0_(*s, s*), giving:

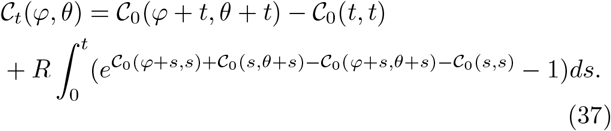

which satisfies the boundary condition, *𝒞*_*t*_(0, 0) = 0 ∀ *t*. Then,

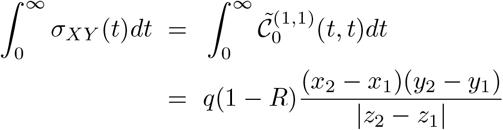

□

If this approximation is accurate, it follows that Proposition 3 also holds for already-recombining populations, but the magnitude of the negative time-integrated covariance (i.e., the magnitude of the average recombinant advantage) decreases as recombination rate increases. This is evidenced by the new factor (1 − *R*) introduced here. However, we do not state this formally, as we have not conducted a thorough analysis and/or exploration of parameter space to determine the accuracy of this approximation.

From Eq (34), the role of recombination in the evolution of total mean fitness, 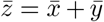, is elucidated by the derivative expressions:

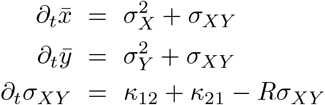

In the absence of selection, the *κ*’s will be zero, giving the prediction that *σ*_*XY*_ (*t*) = *σ*_*XY*_ (0)*e*^−*Rt*^ under neutral evolution. This equation accurately predicts covariance dynamics in simulations of neutral evolution (Fig 7). With selection, the *κ*’s will be non-zero and covariance dynamics are given by:

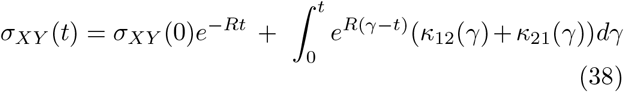

which also shows good agreement with simulations (Fig 8).

**FIG. 7.**
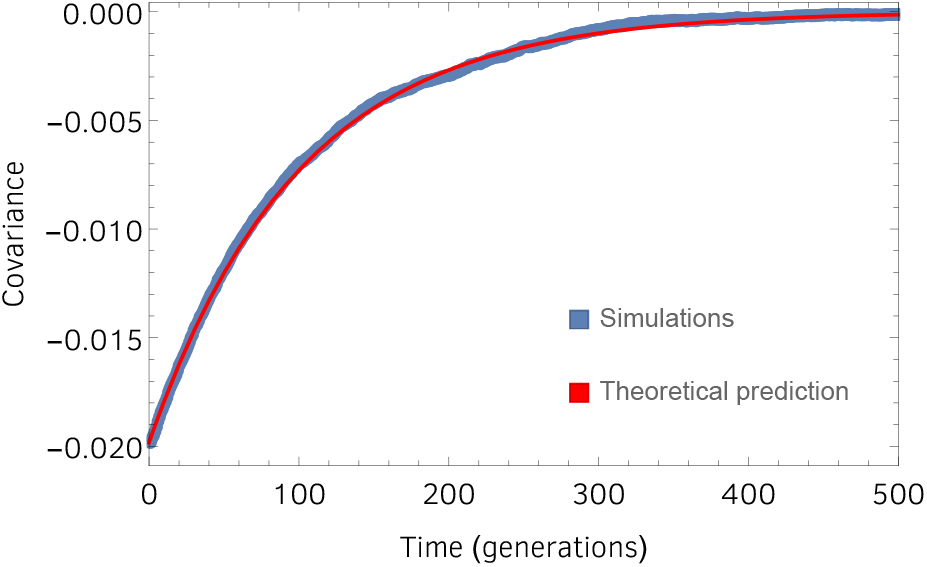
Covariance dynamics under neutral evolution with recombination. Blue dots plot the average of 200 fully stochastic simulations. The red curve plots our theoretical prediction, *σ*_*XY*_ (*t*) = *σ*_*XY*_ (0)*e*^*−Rt*^, which derives from Eq (34). Parameters are: *N* = 10, 000, *R* = 0.01.

**FIG. 8.**
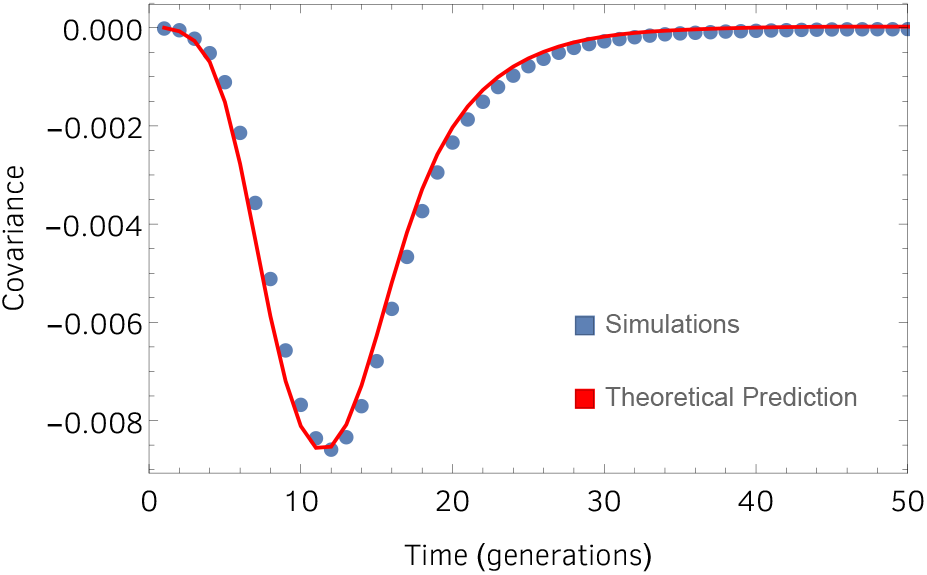
Covariance dynamics under adaptive evolution with recombination. Blue dots plot the average of 200 fully stochastic simulations. The red curve plots the theoretical prediction given by Eq (38). Parameters are: *N* = 10, 000, *R* = 0.01.

### Effect of increasing recombination rate in an already recombining population

Until now, our focus has been on *rec*^+^ modifiers in an otherwise *rec*^−^ population. We now examine the case in which the resident population is *rec*^+^ and the modifier introduced has a higher recombination rate than the resident population. The evolutionary dynamics model developed above allows us to address this question. Specifically, we ask how a further increase in recombination rate increases the rate of increase in mean fitness. Mean fitness increases as 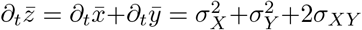, which increases in recombination rate as 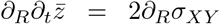 where *σ*_*XY*_ is given by Eq (38). This quantity is positive, as seen in Fig 9.

**FIG. 9.**
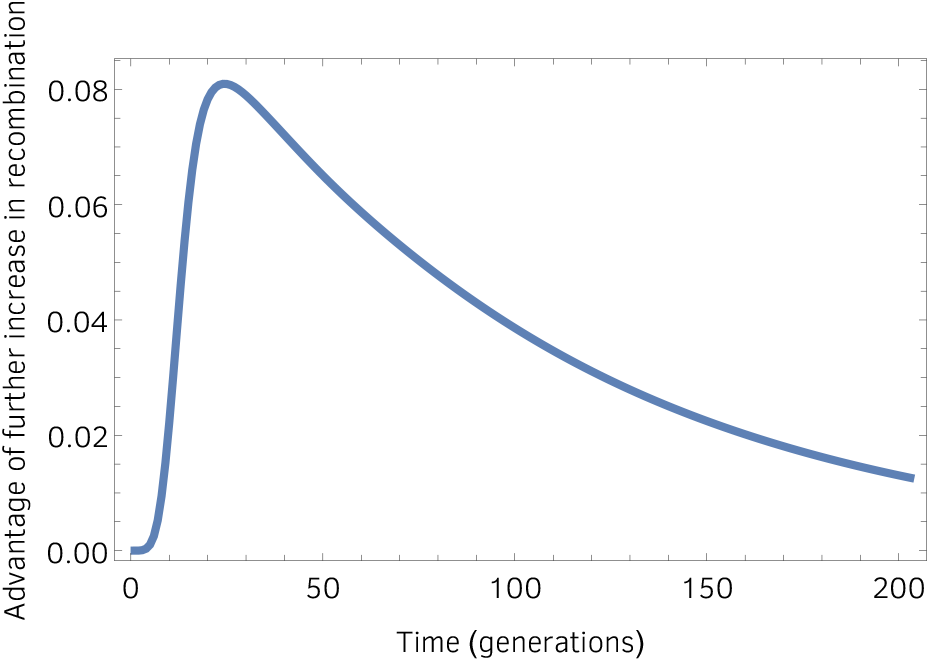
Advantage of further increase in recombination rate in an already-recombining population. This is not selective advantage as it is commonly defined. Instead, it is how fast the rate of fitness increase grows with increase in recombination rate, calculated at the resident recombination rate. Specifically, it is the quantity 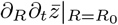, where *R*_0_ is the resident recombination rate. Parameters are: *N* = 10, 000, *R*_0_ = 0.01.

## VI. STATISTICAL MECHANICS OF SEX

We now note a curious connection between the developments presented here and statistical mechanics. This section may serve as a springboard for further work. We believe there is no existing analogue to multilocus evolution in statistical mechanics, in which case these directions may bring something new to both fields.

### Statistical mechanics of single-locus evolution

Based on our developments and our use of the empirical *cgf* (Eq (4)), when there is only one locus under selection, the expected number of individuals with genotype *i* at time *t* may be written as:

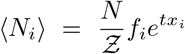

where *N* = total population size, *f*_*i*_ is the initial frequency of genotype *i, x*_*i*_ is the fitness of genotype *i*, and:

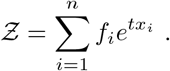

This expression, derived directly from our initial evolution equation, makes the connection between Darwinian evolution and statistical mechanics explicit. We compare these expressions with the Maxwell-Boltzmann equation for number of particles in energy state *ε*_*i*_:

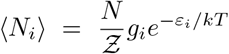

where

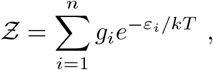

the partition function.

The analogous quantities are shown in Table I. Fitness is analogous to minus the energy. This makes sense, because fitness will tend to increase while energy will tend to decrease. Time is analogous to the inverse temperature. So the evolutionary asymptote as time goes to infinity is analogous to decreasing the temperature to absolute zero. We are not the first to notice a connection between adaptive evolution and statistical mechanics [37–43], but we believe the particular context is new. The multidimensional case to which we now turn does not appear have an immediate analogue in statistical mechanics, yet the extrapolation is straight-forward.

**TABLE I.** A side-by-side comparison of analogous quantities in evolutionary dynamics and statistical mechanics.

| Evolutionary Dynamics | Statistical Mechanics |
| --- | --- |
| fitness, $x_i$ | energy, $-\varepsilon_i$ |
| time, $t$ | inverse temperature, $1/kT$ |
| $t \rightarrow \infty$ | $T \rightarrow 0$ |

### Statistical mechanics of multilocus evolution

Now we extrapolate to multilocus evolution, that is, evolution in which individuals have two or more loci under selection. Here, each genome has *m* loci, each of which contributes to overall fitness *ϕ*. Specifically, genotype *i* has fitness *ϕ*(**x**_*i*_), where vector **x**_*i*_ = (*x*_*i*1_, *x*_*i*2_, …, *x*_*im*_) quantifies fitness contributions from each of the *m* loci. The expected number of individuals with fitness *ϕ*(**x**_*i*_) at time *t* is now:

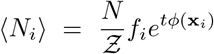

where the partition function is now:

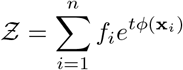

We have found that, as time increases, a population will tend to evolve negative associations among the fitness contributions at the different loci. These negative associations build up across populations [2] and are generated within a single population, in expectation, as time passes. The analogy to statistical mechanics would be that each energy level has some number *m* of contributing factors that determine the energy of that level. As temperature is reduced, negative associations among the contributing factors will be generated. For the system to achieve the lowest possible total energy, the contributing factors would somehow need to be shuffled (analogous to recombination).

## VII. DISCUSSION

### A note about epistasis

Epistasis is non-additivity in genic contributions to fitness, and negative epistasis is concave non-additivity, such that total fitness is always less than the sum of genic fitnesses. A population at equilibrium will harbor negative LD if its constituents exibit negative epistasis across loci. It was once thought that the source of selective imbalance with the strongest causal link to the evolution of sex and recombination was negative, or synergistic, epistasis. However, under negative epistasis, recombinants are initially selectively *suppressed* on average, because recombination breaks up well-matched alleles across loci (it increases diffusion on a concave surface). This initial suppression of recombinants is eventually reversed, and recombinants are eventually amplified by selection, on average, owing to the fact that their fitnesses come from a distribution with larger variance [44]. Whether or not recombinants are ultimately successful therefore depends on the dynamics of recombinant fitness and whether or not the selective reversal is quick enough to rescue and amplify recombinants. Barton [15] identified a small interval of weakly negative epistasis, below zero and not containing it, for which the initial suppression of recombinants was small enough and the selective reversal quick enough to make the recombinants successful on average [3]. As there is no compelling evidence to suspect that epistasis in nature tends to fall within this “goldilocks zone” (or tends to show any general bias away from from zero) [3, 45, 46], epistasis-based explanations for the evolution of sex fell out of favor.

Here, we find that, under natural selection, there is an interval for epistasis outside of which the evolution of recombination would not be favored, but: 1) this interval can be much bigger than that identified under the equilibrium / negative epistasis explanatory framework and, more importantly, 2) it always contains zero (Fig 3). Point 2 is especially relevant because it means that epistasis is not necessary for the evolution of recombination where natural selection is acting. Point 1 suggests that even if a persistent bias in epistasis is demonstrated across the tree of life, it would not invalidate our theory, as long as the bias is not too large.

### Assessment and concluding remarks

Previous studies have found that, when a population undergoing continuous (non-discrete) mutation and recombination achieves equilibrium, a new variant with a lower recombination rate introduced into the population at low frequency will always invade [47–49]. This remarkably encompassing finding was dubbed a “general reduction principle” for recombination [47–49] (also for mutation and migration [47, 50]) and has been interpreted as providing mathematical rigor to the view that the ubiquity of sex and recombination in nature is enigmatic. The discrepancy between the predictions of these models and observations in nature, however, may instead point to the increasingly-accepted view that populations are not at equilibrium most of the time. Analogies between evolutionary genetics and statistical physics are increasingly moving toward non-equilibrium thermodynamics as the more appropriate analogy [51–55].

The setting we analyze in this paper – natural selection simply acting on heritable variation – is one of a system not at equilibrium. Even when our initial fitness distribution is the “mutation-selection-balance distribution” (SM), the fact that no further mutation takes place puts the system out of equilibrium. The very encompassing recombination-augmenting tendency we describe in this paper may perhaps be seen as providing a “general inflation principle” of sorts, for the non-equilibrium case. In light of our findings, recombination is perhaps not the enigma that it has been thought to be. Selection pressure for recombination should appear anywhere natural selection is acting. The evolution of sex and recombination may therefore have evolved less as a catalyst of adaptation and more as a byproduct.

## Acknowledgements

Much of this work was performed during a CNRS-funded visit (P.G.) to the Laboratoire Jean Kuntzmann, University of Grenoble Alpes, France, and during a visit to Bielefeld University (P.G.) funded by Deutsche Forschungsgemeinschaft (German Research Foundation, DFG) via Priority Programme SPP 1590 Probabilistic Structures in Evolution, grants BA 2469/5-2 and WA 967/4-2. P.G. and A.C. received financial support from the USA/Brazil Fulbright scholar program. P.G. and P.S. received financial support from National Aeronautics and Space Administration grant NNA15BB04A. P.G. received further support from the National Institute Of General Medical Sciences of the National Institutes of Health under Award Number R35GM137919 (awarded to Gideon Bradburd). The authors thank S. Otto and N. Barton for their thoughts on early stages of this work. Special thanks go to E. Baake for her thoughts on later stages of this work and help with key mathematical aspects. The authors thank D. Chencha, J. Streelman, R. Rosenzweig and the Biology Department at Georgia Institute of Technology for critical infrastructure and computational support.

## Supplementary Materials

Methods

Supplementary

Text Figs S1 to S12

Tables S1 to S3

References (31 to 40)

## Author contributions

P.G. conceived the theory conceptually; P.G., P.S., B.S. and A.C. developed the theory verbally and with simulation; P.G, B.Y. and J.C. developed the theory mathematically; B.Y. and J.C. provided mathematical proofs for the across-population part; P.G., V.V., F.C. and N.H. provided mathematical proofs for the within-population part. P.G. wrote the paper with critical help and guidance from B.S., P.S. and B.Y.

## Competing interests

The authors declare no competing interests.

## Notes

https://www.biorxiv.org/content/10.1101/2021.06.07.447320v3

https://www.biorxiv.org/content/10.1101/2020.08.28.271486v8

### Competing Interest Statement

The authors have declared no competing interest.

## References

[1] P. J. Gerrish, B. Galeota-Sprung, F. Cordero, P. Sniegowski, A. Colato, N. Hengartner, V. Vejalla, J. Chevallier, and B. Ycart, Natural selection and the advantage of recombination, Phys. Rev. Lett. In Review (2021).

[2] P. J. Gerrish, B. Galeota-Sprung, P. Sniegowski, J. Chevallier, and B. Ycart, Natural selection promotes the evolution of recombination 1: among selected genotypes, Physical Review E In Review (2021).

[3] S. P. Otto and T. Lenormand, enResolving the paradox of sex and recombination, Nat. Rev. Genet. 3, 252 (2002).

[4] N. H. Barton and B. Charlesworth, enWhy sex and recombination?, Science 281, 1986 (1998).

[5] S. P. Otto, enSelective interference and the evolution of sex, J. Hered. 112, 9 (2021).

[6] S. p. Otto, The evolutionary enigma of sex, Am. Nat. 174, S1 (2009).

[7] J. A. G. M. de Visser and S. F. Elena, enThe evolution of sex: empirical insights into the roles of epistasis and drift, Nat. Rev. Genet. 8, 139 (2007).

[8] M. Hartfield and P. D. Keightley, enCurrent hypotheses for the evolution of sex and recombination, Integr. Zool. 7, 192 (2012).

[9] S. C. Lee, M. Ni, W. Li, C. Shertz, and J. Heitman, enThe evolution of sex: a perspective from the fungal kingdom, Microbiol. Mol. Biol. Rev. 74, 298 (2010).

[10] R. A. Fisher, The genetical theory of natural selection (Oxford Clarendon Press, 1930) p. 302.

[11] H. J. Muller, Some genetic aspects of sex, Am. Nat. 66, 118 (1932).

[12] D. Roze and N. H. Barton, enThe Hill-Robertson effect and the evolution of recombination, Genetics 173, 1793 (2006).

[13] W. G. Hill and A. Robertson, enThe effect of linkage on limits to artificial selection, Genet. Res. 8, 269 (1966).

[14] N. H. Barton, enLinkage and the limits to natural selection, Genetics 140, 821 (1995).

[15] N. H. Barton, enA general model for the evolution of recombination, Genet. Res. 65, 123 (1995).

[16] N. H. Barton, enGenetic linkage and natural selection, Philos. Trans. R. Soc. Lond. B Biol. Sci. 365, 2559 (2010).

[17] S. P. Otto and N. H. Barton, enThe evolution of recombination: removing the limits to natural selection, Genetics 147, 879 (1997).

[18] J. F. Crow and M. Kimura, Evolution in sexual and asexual populations, Am. Nat. 99, 439 (1965).

[19] Y. Kim and H. A. Orr, enAdaptation in sexuals vs. asexuals: clonal interference and the Fisher-Muller model, Genetics 171, 1377 (2005).

[20] J. M. Smith and J. Maynard-Smith, The evolution of sex, Vol. 4 (Cambridge University Press Cambridge, 1978).

[21] J. Maynard Smith, The evolution of sex,.

[22] B. Charlesworth, enMutation-selection balance and the evolutionary advantage of sex and recombination, Genet. Res. 89, 451 (1990).

[23] S. P. Otto and M. W. Feldman, enDeleterious mutations, variable epistatic interactions, and the evolution of recombination, Theor. Popul. Biol. 51, 134 (1997).

[24] A. Blachford and A. F. Agrawal, enAssortative mating for fitness and the evolution of recombination, Evolution 60, 1337 (2006).

[25] N. H. Barton and S. P. Otto, enEvolution of recombination due to random drift, Genetics 169, 2353 (2005).

[26] S. P. Otto and N. H. Barton, enSelection for recombination in small populations, Evolution 55, 1921 (2001).

[27] P. D. Keightley and S. P. Otto, enInterference among deleterious mutations favours sex and recombination in finite populations, Nature 443, 89 (2006).

[28] S. P. Otto, enThe evolutionary enigma of sex, Am. Nat. 174 Suppl 1, S1 (2009).

[29] G. Martin and L. Roques, enThe nonstationary dynamics of fitness distributions: Asexual model with epistasis and standing variation, Genetics 204, 1541 (2016).

[30] M.-E. Gil, F. Hamel, G. Martin, and L. Roques, Mathematical properties of a class of integro-differential models from population genetics, SIAM J. Appl. Math. 77, 1536 (2017).

[31] Y. Anciaux, A. Lambert, O. Ronce, L. Roques, and G. Martin, enPopulation persistence under high mutation rate: From evolutionary rescue to lethal mutagenesis, Evolution 73, 1517 (2019).

[32] P. J. Gerrish and P. D. Sniegowski, enReal time forecasting of near-future evolution, J. R. Soc. Interface 9, 2268 (2012).

[33] R. Bürger, enMoments, cumulants, and polygenic dynamics, J. Math. Biol. 30, 199 (1991).

[34] M. Smerlak and A. Youssef, enLimiting fitness distributions in evolutionary dynamics, J. Theor. Biol. 416, 68 (2017).

[35] J. Felsenstein and S. Yokoyama, enThe evolutionary advantage of recombination. II. individual selection for recombination, Genetics 83, 845 (1976).

[36] S. P. Otto and T. Lenormand, Resolving the paradox of sex and recombination, Nat. Rev. Genet. 3, 252 (2002).

[37] N. H. Barton and H. P. de Vladar, Statistical mechanics and the evolution of polygenic quantitative traits (2009).

[38] M. Smerlak, Natural selection as coarsening, J. Stat. Phys. (2018).

[39] D. Fisher, M. Lässig, and B. Shraiman, enEvolutionary dynamics and statistical physics, J. Stat. Mech. 2013, N01001 (2013).

[40] E. Baake and H. Wagner, Mutation–selection models solved exactly with methods of statistical mechanics, Genet. Res. 78, 93 (2001).

[41] de Vladar Harold P. and Barton Nick H., The statistical mechanics of a polygenic character under stabilizing selection, mutation and drift, J. R. Soc. Interface 8, 720 (2011).

[42] H. P. de Vladar and N. H. Barton, enThe contribution of statistical physics to evolutionary biology, Trends Ecol. Evol. 26, 424 (2011).

[43] T. C. B. McLeish, enAre there ergodic limits to evolution? ergodic exploration of genome space and convergence, Interface Focus 5, 20150041 (2015).

[44] B. Charlesworth and N. H. Barton, Recombination load associated with selection for increased recombination (1996).

[45] S. P. Otto, enUnravelling gene interactions, Nature 390, 343 (1997).

[46] W. R. Rice, enExperimental tests of the adaptive significance of sexual recombination, Nat. Rev. Genet. 3, 241 (2002).

[47] L. Altenberg, U. Liberman, and M. W. Feldman, Unified reduction principle for the evolution of mutation, migration, and recombination, Proceedings of the National Academy of Sciences 114, E2392 (2017).

[48] U. Liberman and M. W. Feldman, enA general reduction principle for genetic modifiers of recombination, Theor. Popul. Biol. 30, 341 (1986).

[49] M. W. Feldman and U. Liberman, An evolutionary reduction principle for genetic modifiers, Proceedings of the National Academy of Sciences 83, 4824 (1986).

[50] U. Liberman and M. W. Feldman, Modifiers of mutation rate: A general reduction principle, Theor. Popul. Biol. 30, 125 (1986).

[51] V. Mustonen and M. Lässig, enFitness flux and ubiquity of adaptive evolution, Proc. Natl. Acad. Sci. U. S. A. 107, 4248 (2010).

[52] C. Payen, A. B. Sunshine, G. T. Ong, J. L. Pogachar, W. Zhao, and M. J. Dunham, enHigh-Throughput identification of adaptive mutations in experimentally evolved yeast populations, PLoS Genet. 12, e1006339 (2016).

[53] D. W. Hall and S. B. Joseph, enA high frequency of beneficial mutations across multiple fitness components in saccharomyces cerevisiae, Genetics 185, 1397 (2010).

[54] I. Gordo, enEvolutionary change in the human gut microbiome: From a static to a dynamic view, PLoS Biol. 17, e3000126 (2019).

[55] L. Perfeito, L. Fernandes, C. Mota, and I. Gordo, enAdaptive mutations in bacteria: high rate and small effects, Science 317, 813 (2007).

